# Angiotensin II induces apoptosis in human induced pluripotent stem cell-derived cardiomyocytes

**DOI:** 10.1101/2020.03.19.998344

**Authors:** Qiang Gao, Ping Wang, Zhiming Wu, Hailong Qiu, Bin Lin, Jimei Chen, Jianzheng Cen, Jian Zhuang

**Affiliations:** Department of Cardiac Surgery, Guangdong Cardiovascular Institute, Guangdong Provincial People’s Hospital, Guangdong Academy of Medical Sciences, Guangzhou, Guangdong 510100, China; School of Medical Imaging, Tianjin Medical University, Tianjin 300203, China; Department of Urology, Sun Yat-sen University Cancer Center, State Key Laboratory of Oncology in South China, Collaborative Innovation Center for Cancer Medicine, Guangzhou 510060, China; Guangdong Beating Origin Regenerative Medicine Co. Ltd., Foshan, Guangdong 528231, China

**Keywords:** Angiotensin II, concentration, iPSC-CM, apoptosis, cell viability

## Abstract

**Background:** The renin-angiotensin system (RAS) functions fundamentally to regulate the pathological process of cardiovascular diseases, such as heart failure and hypertension. As the major effector in RAS, angiotensin II activates angiotensin II receptors to initiate the downstream pathways, which lead to the phenotypes including apoptosis, hypertrophy, and cardiac remodeling. Human induced pluripotent stem cell-derived cardiomyocytes (iPSC-CM) are being applied as a promising platform for personalized medicine to heart diseases. However, whether angiotensin II induces apoptosis in iPSC-CM is still obscure, which raises an uncertainty about the clinical applications of iPSC-CM.

**Methods:** We treated iPSC-CM with angiotensin II at eight concentrations (0 nM, 1 nM, 10 nM, 100 nM, 1 μM, 10 μM, 100 μM and 1 mM) and four incubation durations (24 hours, 48 hours, 6 days and 10 days), then PrestoBlue reagent and a apoptosis marker were used to examine the viability and apoptosis status of cardiomyocytes from each group. The expression levels of some apoptosis and proliferation related genes were also analyzed.

**Results:** High concentration angiotensin II with a long-term treatment caused apoptosis and cell viability drop-off in iPSC-CM. Specifically, under a 10-day treatment with 1 mM angiotensin II, the viability of iPSC-CM was reduced by an average of 41% (*p*=2.073E-08), and the percentage of apoptotic cells was 2.74 times higher than the controls averagely (*p*=6.248E-12). The data mining of previous RNA-seq data revealed that angiotensin II receptor type I was the major receptor in iPSC-CM. Conclusions: For the first time, our data confirmed the apoptotic effect of angiotensin II to iPSC-CM. The angiotensin II concentrations and exposure time for apoptosis induction were depicted in our study, which provided supports to iPSC-CM as an *in vitro* model for cardiovascular disease study.

## Introduction

Heart failure (HF) is the most popular chronic disease among the elderly in developed countries (1), and its prevalence and incidence is increasing in the resent years (2-4). Approximately 26 million individuals are suffering HF worldwide (5), in which the survival rate within 5 years of admission was less than 53% (6-8). The chronic heart failure causes a tremendous economic burden with an estimation of $108 billion per year globally (9). As a comprehensive medical condition, HF is an end stage of many cardiovascular diseases, such as ischaemic heart disease, chronic obstructive pulmonary disease, hypertensive heart disease, and rheumatic heart disease (10).

Like other multifactorial conditions, progression of HF is modified by the genetic diversity of affected individuals, which makes routine HF treatment difficult to achieve satisfying benefits (4,11). It is an urgent mission to establish an *in vitro* system to evaluate the effects of different medicine to each HF patient. The advances of cell reprogramming technology (especially induced pluripotent stem cell (iPSC) technology) (12,13), cardiomyocyte differentiation (14,15) and purification (16,17) protocols provide us powerful tools to generate cardiomyocytes from patients’ somatic cells. These iPSC-derived cardiomyocytes (iPSC-CM) could mimic the cardiac phenotypes from the donors without ethics issues. To date, iPSC-CM were used to investigate long QT syndrome (18-20), arrhythmia (21,22), hypertrophic cardiomyopathy (23), dilated cardiomyopathy (24,25), mitochondrial cardiomyopathy (26), Duchenne muscular dystrophy (27), catecholaminergic polymorphic ventricular tachycardia (28), and other cardiovascular diseases.

On the other side, due to the high-throughput advantages, iPSC-CM were more and more applied to pharmacologic and cardiotoxicity testing (29-35). There medications, including aldosterone antagonist, angiotensin-converting enzyme inhibitors, angiotensin II receptor blockers, angiotensin receptor-neprilysin inhibitors, and beta-blockers (36,37), are available for testing in the patients’ iPSC-CM. Notably, many of these drugs target to the angiotensin pathway. Indeed, angiotensin II (Ang II) plays a crucial role in the pathological mechanism of HF (38-40). It was reported that Ang II caused apoptosis of cardiomyocytes in rats (41-43), vascular fibrosis and damage (44). In fact, Ang II was an apoptosis inducer of many kinds of cells, such as vascular smooth muscle cells (45), human endothelial cells (46), coronary artery endothelial cells (47), rat glomerular epithelial cells (48), alveolar epithelial cells (49), intestinal epithelial cells (50), renal proximal tubular cells (51,52), mesangial cells (53,54), and podocytes (55). However, there are no studies reporting that Ang II induced apoptosis in iPSC-CM. Instead, several reports showed that Ang II could induce hypertrophic phenotypes in iPSC-CM (56,57). Since the response to Ang II is a critical criterion for drug testing, it is necessary to figure out whether Ang II induces apoptosis in iPSC-CM.

In this research, we performed short-term and long-term treatments of Ang II to iPSC-CM. We presented that only long-term treatments with high concentration Ang II caused apoptosis in iPSC-CM, and Ang II triggered the downstream signaling cascades mainly through angiotensin II receptor type 1 (AT1).

## Results

### Cell viability was significantly reduced in iPSC-CM under long-term treatments of high concentration Ang II

To determine the pro-apoptotic effects by Ang II at various concentrations and incubation time, 8 concentrations (0 nM (untreated), 1 nM, 10 nM, 100 nM, 1 μM, 10 μM, 100 μM and 1 mM) and 4 treatment durations (24 hours, 48 hours, 6 days and 10 days) were tested in iPSC-CM. Here, 1 nM to 1 μM were defined as “low concentrations”, while 10 μM, 100 μM and 1 mM were defined as “high concentrations”; similarly, 24 hours and 48 hours were defined as “short terms”, while 6 days and 10 days were defined as “long terms”. Human iPS cell line DYR0100 (ATCC) was differentiated to cardiomyocytes by using the protocols from previous reports (14,16). After metabolic selections (16), iPSC-CM with > 98% purity were seeded to 96-well plates for recovery. On differentiation day 30, iPSC-CM were applied to Ang II treatments. Under short-term treatments (24 hours and 48 hours) of Ang II, there were no significant differences between control iPSC-CM and Ang II-treated iPSC-CM in cell viability, no matter with low or high concentrations (Figure 1a). Conversely, after 6-day treatments, 100 μM and 1 mM Ang II significantly reduced the viability of iPSC-CM (Fold change of cell viability, 100 μM vs. untreated control: 0.763, *p*=8.085E-07; 1 mM vs. untreated control: 0.762, *p*=7.306E-07); after 10-day treatments, 10 μM, 100 μM and 1 mM Ang II caused ∼21%, 36% and 41% drop-off in cell viability, respectively (Fold change of cell viability, 10 μM vs. untreated control: 0.795, *p*=0.034; 100 μM vs. untreated control: 0.638, *p*=4.885E-07; 1 mM vs. untreated control: 0.590, *p*=2.073E-08). Ang II at other concentrations seemed to slightly decrease the cell viability without statistical significance (Figure 1b). Intriguingly, we observed that iPSC-CM under Ang II treatment for a short term showed a light increase in viability, which was inconsistent with the data from other types of cells.

**Figure 1.**
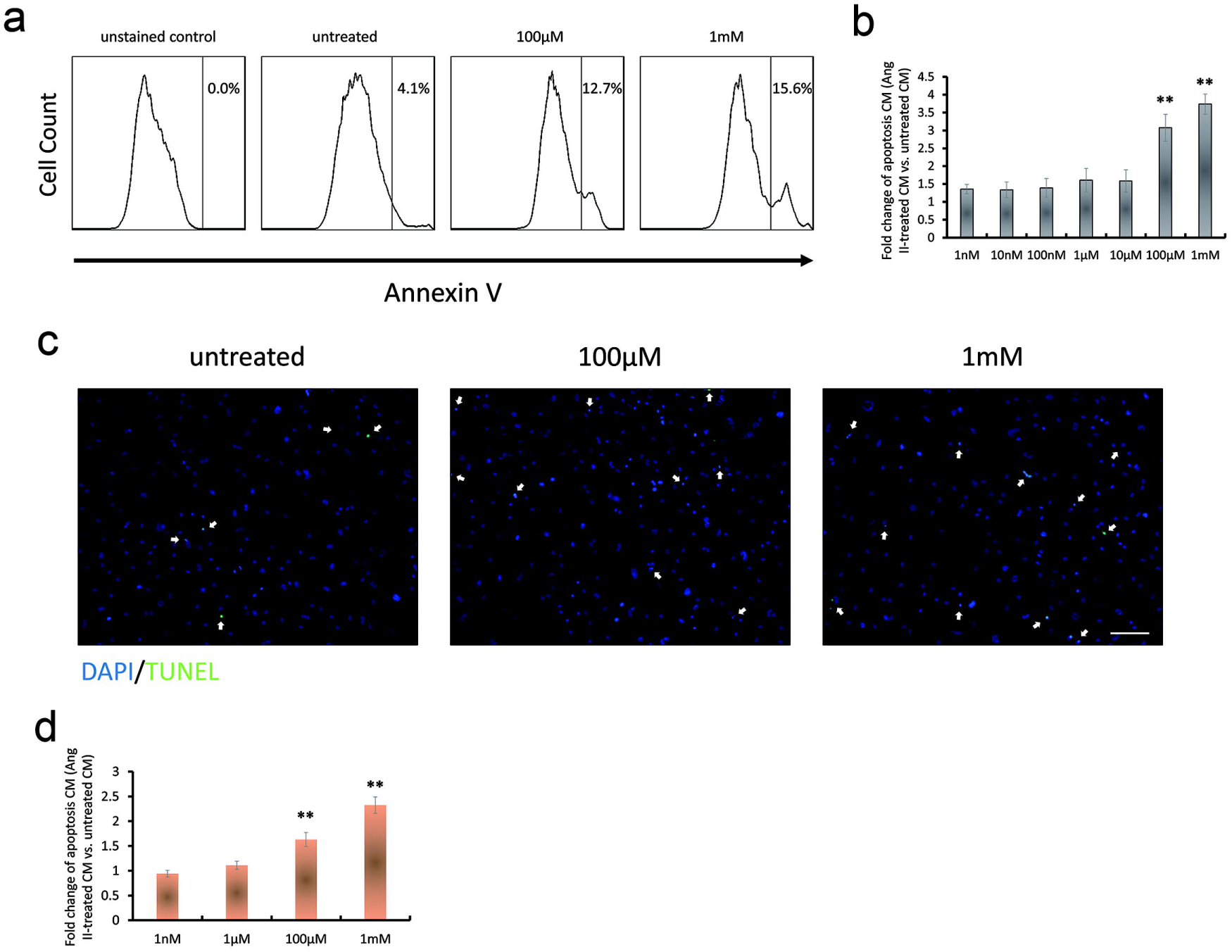
Cell viability was decreased after long-term Ang II treatment. The effect of a series of Ang II concentrations (1 nM, 10 nM, 100 nM, 1 μM, 10 μM, 100 μM, 1 mM) to iPSC-CM was evaluated by PrestoBlue reagent. The relative fluorescence unit (RFU) fold changes from untreated cardiomyocytes to Ang II-treated cardiomyocytes were calculated in a), short-term (24 hours and 48 hours) treatment set and b), long-term (6 days and 10 days) treatment set (n=7). #, *p*<0.05; **, *p*<0.001; unlabeled data, no significant differences.

### Long-term incubation of high concentration Ang II induced apoptosis in iPSC-CM

Based on the results of cell viability test, the viability of iPSC-CM only declined under long-term treatments of Ang II. Therefore, to investigate the apoptotic rates, we performed apoptosis analysis in iPSC-CM after 10-day treatments of Ang II. iPSC-CM under 10-day incubations were collected and stained by Annexin V conjugated with Alexa Fluor 488, then analyzed by fluorescence-activated cell sorting (FACS). As Figure 2 shown, the apoptotic rate of untreated iPSC-CM was about 4.1%, whereas about 12.7% and 15.6% of iPSC-CM were apoptotic after 100 μM and 1 mM Ang II treatments, respectively (Figure 2a), which indicated that the apoptotic cells under 100 μM Ang II treatment were averagely 3.08 times as many as untreated group (*p*=2.029E-09), while the apoptotic cells under 1 mM Ang II treatment were averagely 3.74 times as many as untreated group (*p*=6.248E-12) (Figure 2b). There were no significant differences between treatments with other concentrations and controls. TUNEL assay was also performed to verify the apoptosis analysis. iPSC-CM were treated with 1 nM, 1 μM, 100 μM and 1 mM Ang II for 10 days then applied to TUNEL assay. The TUNEL positive cells were significantly increased in 100 μM and 1 mM Ang II-treated iPSC-CM. Compared to the untreated iPSC-CM, 100 μM and 1 mM Ang II-treated iPSC-CM exhibited 1.63 (*p*=7.144E-08) and 2.32-fold (*p*=1.298E-15) increase in apoptotic rate, respectively (Figure 2c, d). 1 nM and 1 μM Ang II treatments did not show differences compared with the control set.

**Figure 2.**
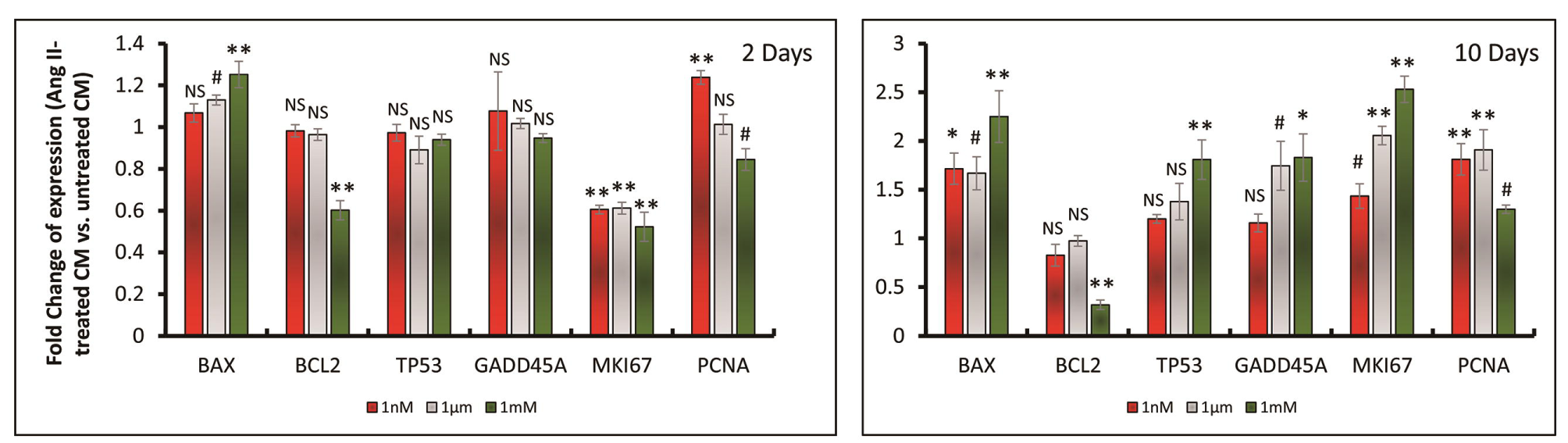
Long-term Ang II treatment caused apoptosis in iPSC-CM. a) Representative fluorescence-activated cell sorting (FACS) analyses of apoptosis marker Annexin V in iPSC-CM with 0 nM (untreated), 100 μM and 1 mM Ang II treatment for 10 days. In the unstained group, the DPBS buffer was used instead of the Annexin V staining reagent. b) The proportion of Annexin V-positive cells (PAP) represented the apoptosis status of iPSC-CM. The fold changes of PAP from untreated cardiomyocytes to Ang II-treated cardiomyocytes at different concentrations were calculated (n=3). **, *p*<0.001; unlabeled data, no significant differences. c) Representative images of TUNEL assay in iPSC-CM with 0 nM (untreated), 100 μM and 1 mM Ang II treatment for 10 days. Nuclei were stained with DAPI (blue). The white arrows denote TUNEL-positive cardiomyocyte nuclei (green). The scale bar is 100 μm. d) The proportion of TUNEL-positive cells (TP) also reflected the apoptosis status of iPSC-CM. The fold changes of TP from untreated cardiomyocytes to Ang II-treated cardiomyocytes at 0 nM (untreated), 1 nM, 1μM, 100 μM and 1 mM concentrations were calculated. Approximately 5,000 cells were counted in each group. **, *p*<0.001; unlabeled data, no significant differences.

We next evaluated the expression levels of apoptosis and proliferation related genes in iPSC-CM, which were treated with Ang II at different concentrations (1 nM, 1 μM and 1 mM) for 2 days and 10 days (Figure 3). In the iPSC-CM under a 2-day Ang II treatment, 1 nM and 1 μM Ang II did not change the expression levels of *BCL2, TP53* and *GADD45A*, while 1 mM Ang II led to 1.25-fold increase in *BAX* (*p*=0.0004) and down-regulation of *BCL2* (*p*=1.432E-06). The expression of *MKI67* was decreased after Ang II treatment at all the three concentrations. On the other hand, after the 10-day incubation with 1 mM Ang II, the expression of *BAX, TP53, GADD45A* and *MKI67* was significantly up-regulated (with 2.25-, 1.81-, 1.83-, and 2.53-fold increase, respectively; P values were 0.0001, 0.0006, 0.0030, 5.348E-07, respectively), while the expression of *BCL2* dropped by 70% (*p*=8.583E-06). Contrary to the 2-day treatment, 10-day treatment elevated the expression of *MKI67* at all the three concentrations. Encoding a pro-apoptotic protein, *BAX* functions as an apoptotic activator in the apoptosis pathway (58). In contrast, BCL2 was an anti-apoptotic molecule that promoted cell survival (59) and blocked programmed cell death (60). *TP53* and *GADD45A* were cell cycle-regulated genes, which were also reported to mediate apoptosis in cells (61,62). *MKI67* and *PCNA* were markers of cell proliferation (63). It is worth noting that 1 mM Ang II already caused up-regulation of *BAX* and down-regulation of *BCL2* in a short-term treatment, and with the increase of incubation time, the BAX/BCL2 ratio, which was considered as an apoptotic index, was elevated sharply. Besides, 1 mM Ang II increased the expression of *TP53* and *GADD45A* as well, indicating that the cardiomyocytes were in the apoptotic status.

**Figure 3.**
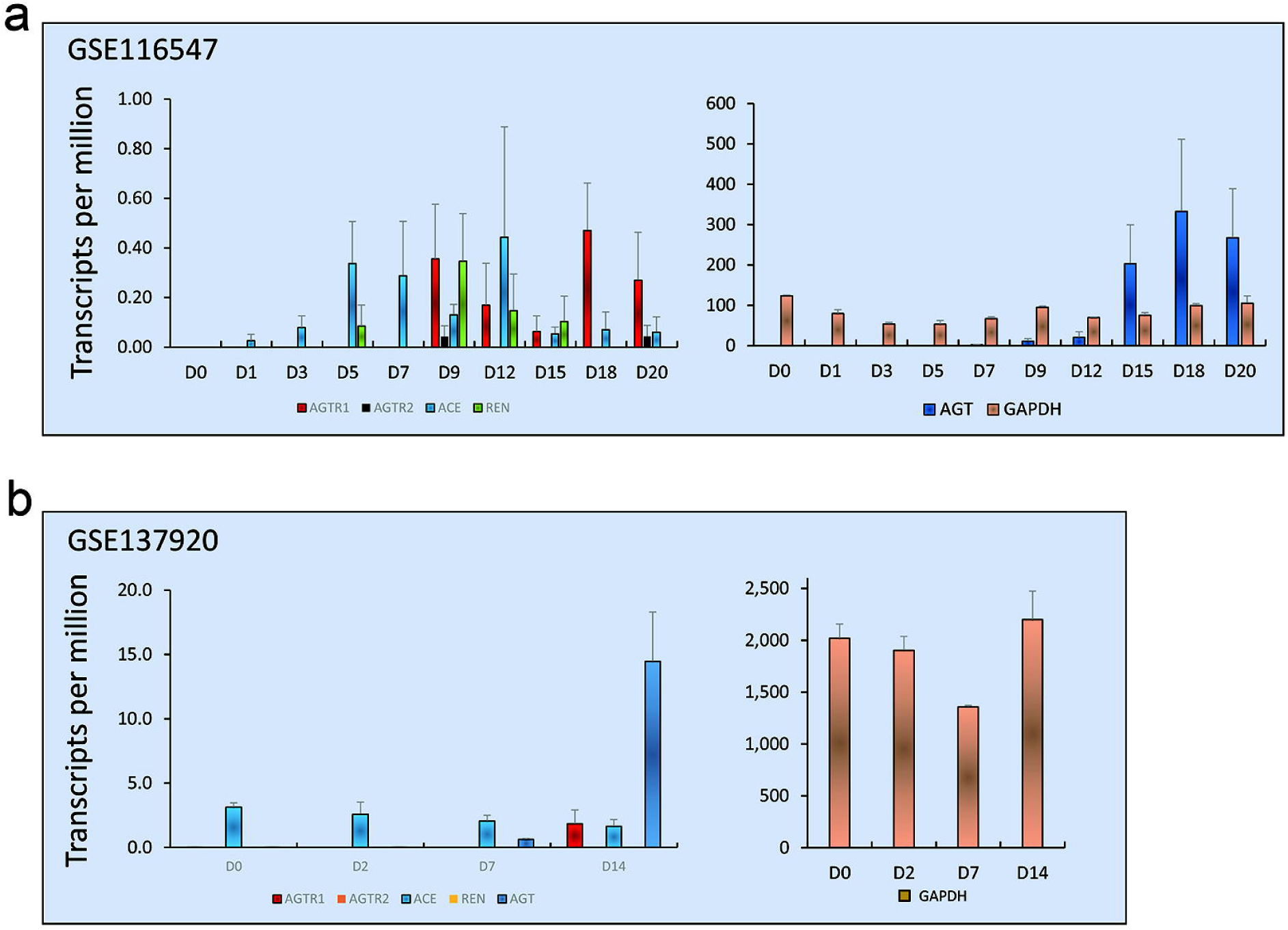
High concentration Ang II was associated with the expression changes of apoptosis and proliferation related genes. iPSC-CM were treated with 0 nM (untreated), 1 nM, 1μM and 1 mM Ang II for 2 days and 10 days. Total mRNA was extracted from each group (n=3). Expression of apoptosis and proliferation related genes in 2-day and 10-day treatments was shown in left and right panels, respectively. #, *p*<0.05; *, *p*<0.01; **, *p*<0.001; NS, no significant differences.

### AT1, not AT2 was expressed in iPSC-CM

There are two types of angiotensin II receptors: AT1 and AT2. Basically, AT1 was implicated in the major effects of Ang II, such as proliferation, hypertrophy, and fibrosis, while AT2 acted to antagonize the effects of AT1 (64,65). Generally, it was considered that Ang II caused apoptosis and cardiac remodeling through binding to AT2 (66,67), however, it was reported that AT1 was also associated with Ang II-mediated apoptosis (68-70). To investigate which receptor was involved in the Ang II-mediated apoptosis in iPSC-CM, we first examined the expression levels of the renin-angiotensin system (RAS)-related genes (*AGTR1, AGTR2, ACE, REN*, and *AGT*) by re-analyzing the expression profiling data of cells from different time points of cardiac differentiation (GSE116574 and GSE137920) (71,72). In the data set GSE116574, *AGTR1* started to be expressed from differentiation day 9, whereas the transcripts of *AGTR2* were too low to be considered undetectable (Figure 4a). In the data set GSE137920, the transcripts of *AGTR1* could be detected on differentiation day 14, however, *AGTR2* was not expressed in the cells from all the time points (Figure 4b). We performed reverse transcription PCR to detect the expression levels of *AGTR1* and *AGTR2* in iPSC-CM on differentiation day 30. In agreement with the previous reports, we found that *AGTR1* was expressed in the iPSC-CM, while *AGTR2* was hardly detectable (data not shown). According to these observations, we proposed that Ang II induced apoptosis in the iPSC-CM through binding to AT1.

**Figure 4.**
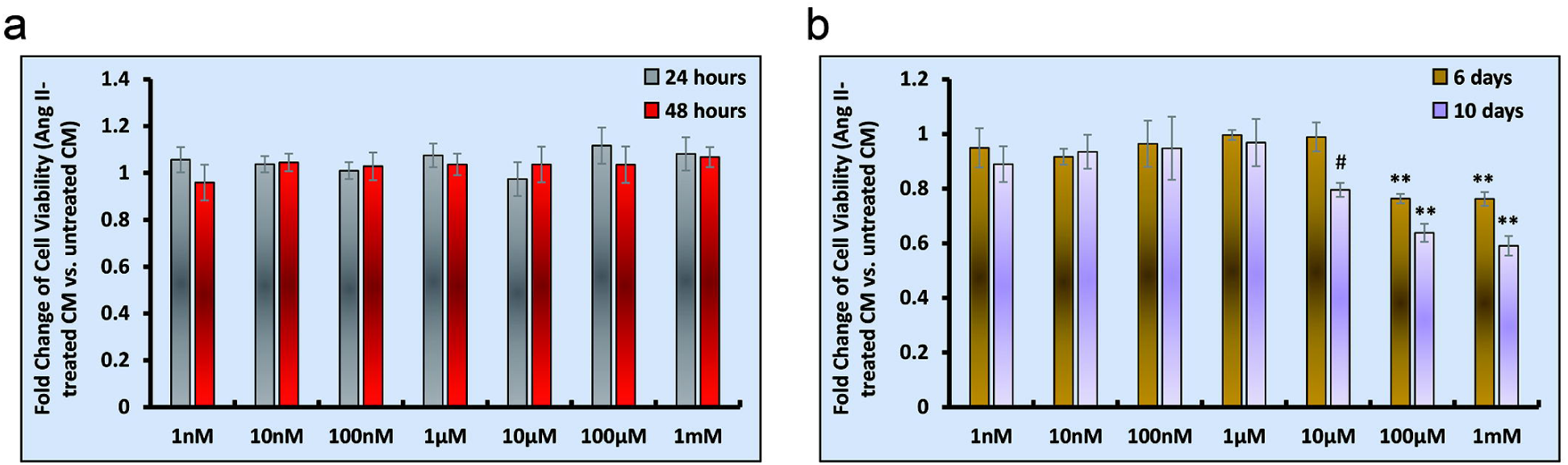
AT1 was the major angiotensin II receptor in iPSC-CM. RNA-seq data about *AGTR1, AGTR2, ACE, REN, AGT* and *GAPDH* genes from a) GSE116547 dataset and b) GSE137920 dataset were re-analyzed. Transcripts per million (TPM) were used to evaluate the expression levels of each gene. D+number denoted the days during cardiomyocyte differentiation.

## Discussion

The renin-angiotensin system is the key pathway engaged in pathogenesis of many cardiovascular diseases, including cardiomyopathy, heart failure and hypertension (73-75). As the central component of RAS, Ang II activated AT1 and AT2 to regulate the physiological functions of cardiomyocytes. Many chemicals targeting RAS, such as angiotensin receptor blockers (valsartan, losartan, etc.), angiotensin converting enzyme inhibitors (captopril, benazepril, enalapril, etc.) and angiotensin receptor-neprilysin inhibitors (Entresto, etc.), were used as medications for cardiovascular disease therapy. Consequently, the response to Ang II was an essential requirement for a cardiovascular disease model. To the best of our knowledge, only a few studies investigated the physiological effects of Ang II to iPSC-CM. Binah group examined the effects of Ang II on the contraction and intracellular Ca^2+^ transient in hESC-CM (76,77). Harding group and Fukuda group reported that Ang II could induce hypertrophy in iPSC and hESC-derived cardiomyocytes (56,78), while Harding group presented a contrary result that Ang II had no significant effect on cell size in hESC-CM and hiPSC-CM three years later (79). Radisic group used 200 nM Ang II to treat hESC-CM in 3D culture for 7 days, and found no significant differences in cell death and viability in drug-treated and untreated hESC-CM (80). This was the first attempt to study the effects after chronic Ang II exposure. It was unexpected that there were no reports demonstrating the apoptotic effects of Ang II to iPSC-CM. For the first time, we showed evidences that only high concentration Ang II (10 μM, 100 μM and 1 mM) provoked apoptosis after chronic treatment, namely, the apoptosis phenotype was dependent on the concentration and incubation time of Ang II.

The expression of AT1 in iPSC-CM was confirmed by immunofluorescent staining (76). However, according to the RNA-seq data (71,72), the expression level of AT1 was relatively very low compared to GAPDH in iPSC-CM (Figure 4). The low expression level of AT1 might explain short-term stimuli of Ang II could not induce apoptosis: when the expression of AT2 was missing, even the high concentration Ang II could not trigger the apoptosis pathway through a small amount of AT1. Since Ang II infusion for 28 days could increase the protein levels of AT1 and AT2 in rats (35% and 100% increase, respectively) (81), we assumed that a long-term Ang II incubation could also up-regulate the expression level of AT2 in iPSC-CM, which offered a reasonable explanation for the apoptosis phenotype induced by long-term Ang II treatments: chronic exposure by high concentration Ang II facilitated iPSC-CM to produce enough AT2 so that Ang II could activate AT2 to initiate downstream apoptosis signaling. In conclusion, we proved that Ang II could induce apoptosis in iPSC-CM in a concentration and time-dependent manner. These findings provided necessary supports for iPSC-CM as a powerful tool to investigate cardiovascular diseases *in vitro*.

## Method

### Cell Culture

Human induced pluripotent stem cells (iPSC) DYR0100 (The American Type Culture Collection, ATCC) were plated on Matrigel matrix (hESC-Qualified, LDEV-Free, Corning, 354277)-coated plates, and then were cultured with StemFlex Medium (Gibco, A3349401). StemFlex Medium was changed every two days. iPSC were passaged every three days or when the cell culture was 80-90% confluent. During passages, iPSC were rinsed with 1X DPBS (Gibco, 14040133) for one time then were treated with 0.5mM EDTA (Invitrogen, 15575020) in 1X DPBS (Gibco, 14190144) for 10 mins at room temperature. The split ratio was 1:3-1:6. The detailed differentiation protocol was described in the previous published reports (16,82). Briefly, iPSC were treated with small molecule CHIR99021 (Tocris, 4423, final concentration 10 μM) in the RPMI-BSA medium [RPMI 1640 Medium (HyClone, SH30027.01) supplemented with 213 μg/ml AA2P (l-ascorbic acid 2-phosphate magnesium) (A8960, Sigma) and 0.1% bovine serum albumin (BSA) (A1470, Sigma)] for 24 hours, then were incubated with RPMI-BSA medium for 48 hours. On differentiation day 4, cells were treated with the small molecule IWP2 (Tocris, 3533, final concentration 5 μM) in RPMI-BSA medium. After 48 hours, medium was changed to RPMI-BSA medium. Then, RPMI 1640 Medium supplemented with 3% KnockOut Serum Replacement (Gibco, 10828-028) was used to culture the cardiomyocytes in the following experiments. A metabolic selection method was used to purify iPSC-derived cardiomyocytes (16). The metabolic selection medium was prepared as DMEM Medium (No Glucose) (Gibco, 11966-025) supplemented with 0.1% BSA (Sigma, A1470) and 1× Linoleic Acid-Oleic Acid-Albumin (Sigma, L9655). Cells were treated with metabolic selection medium for 3-6 days. The medium was changed every 2 days. The purity of cardiomyocytes could reach as high as 99%. After purification, the cardiomyocyte culture medium supplemented with various amount of Ang II (MedChemExpress, HY-13948) was used for Ang II treatment, and the Ang II medium was changed every 2 days.

### Cell Viability Assay

PrestoBlue Cell Viability Reagent (Invitrogen, A13261) was used to detect the viability of cardiomyocytes. PrestoBlue was warmed to room temperature before use. One-tenth volume of PrestoBlue was added to the cardiomyocyte medium to prepare reaction medium. For a 96-well plate, 100 µl reaction media were added to each well. The same amount reaction medium was added to wells without cells as blank controls. Plates were incubated in the 37°C incubator for 60 mins then fluorescence (excitation wavelength: 560 nm, emission wavelength: 590 nm) was read by Varioskan Flash Multimode Reader (Thermo Scientific). The results were normalized by cell numbers.

### Apoptosis Detection Assay

The apoptosis detection kit Annexin V, Alexa Fluor™ 488 conjugate (Invitrogen, V13201) was used to detect the apoptosis status of Ang II-treated and untreated cardiomyocytes. Cardiomyocytes were labeled with apoptosis marker Annexin V then sorted by the FACSAria™ II analyzer (BD). FACS results were analyzed by using FlowJo software.

### TUNEL Assay

TUNEL assay was performed by using TdT In Situ Apoptosis Detection Kit (R&D Systems, 4812-30-K). Cardiomyocytes were dissociated and re-plated on the glass coverslips (Fisher Scientific, S175223). We followed the manufacturer’s protocol except fixed cells were labeled with DAPI before mounted by fluorescence mounting media (Agilent, S302380-2). Images were taken by the DMi6000 B inverted microscope (Leica) and analyzed by using ImageJ software.

### Quantitative PCR

Total RNA was extracted using the UNlQ-10 Column Trizol Total RNA Isolation Kit (Sangon Biotech, B511321-0100) prior to the treatment with DNase I (Sangon Biotech, B618252) for 30 min. mRNA was reverse transcribed using iScript Reverse Transcription Supermix (Bio-Rad, 1708841). Quantitative PCR was performed using a PikoReal Real-Time PCR System (Thermo Fisher) with SsoAdvanced^™^ Universal SYBR^®^ Green Supermix (Bio-Rad, 1725271). The quantitative PCR primers from the previous report (83) are as followed (from 5′ to 3′):

*BAX*-RT-F: CAAACTGGTGCTCAAGGCCC;

*BAX*-RT-R: GGGCGTCCCAAAGTAGGAGA;

*BCL2*-RT-F: CTGGTGGACAACATCGCCCT;

*BCL2*-RT-R: TCTTCAGAGACAGCCAGGAGAAAT;

*MKI67*-RT-F: TGTGCCTGCTCGACCCTACA;

*MKI67*-RT-R: TGAAATAGCGATGTGACATGTGCT;

*PCNA*-RT-F: TTTGGTGCAGCTCACCCTG;

*PCNA*-RT-R: CGCGTTATCTTCGGCCCTTA;

*TP53*-RT-F: GCGTGTTTGTGCCTGTCCTG;

*TP53*-RT-R: TGGTTTCTTCTTTGGCTGGG;

*GADD45A*-RT-F: GATGCCCTGGAGGAAGTGCT;

*GADD45A*-RT-R: GAGCCACATCTCTGTCGTCGT;

*GAPDH*-RT-F: TGGGTGTGAACCATGAGAAG;

*GAPDH*-RT-R: GTGTCGCTGTTGAAGTCAGA.

RNA-seq data re-analysis

Raw fastq files were downloaded from GEO (GSE116574 and GSE137920) by SRA-Tools (https://github.com/ncbi/sra-tools). Low quality bases and adapters were trimmed by trim_galore (http://www.bioinformatics.babraham.ac.uk/projects/trim_galore/). We employed Kallisto (v0.46.0) (84) to quantified transcript abundance as transcripts per million (TPM), using gene annotation in the GENCODE database (v32) (85). Then we summed TPM of all alternative splicing transcripts of a gene to obtain gene expression levels.

### Statistic

Values are expressed as mean ± SD (standard deviation). Statistical significances were evaluated using one-way ANOVA with Bonferroni correction. *P*<0.05 was considered statistically significant.

## Author contributions

(I) Conception and design: Jian Zhuang and Jianzheng Cen

(II) Administrative support: Jian Zhuang, Jianzheng Cen and Jimei Chen

(III) Provision of study materials or patients: Qiang Gao and Lin Bin

(IV) Collection and assembly of data: Qiang Gao, Ping Wang, Zhiming Wu and Hailong Qiu

(V) Data analysis and interpretation: Qiang Gao

(VI) Manuscript writing: All authors

(VII) Final approval of manuscript: All authors

